# Type III-B CRISPR-Cas signaling-based cascade of proteolytic cleavages

**DOI:** 10.1101/2023.06.23.546230

**Authors:** Jurre A. Steens, Jack P.K. Bravo, Carl Raymund P. Salazar, Caglar Yildiz, Afonso M. Amieiro, Stephan Köstlbacher, Stijn H.P. Prinsen, Constantinos Patinios, Andreas Bardis, Arjan Barendregt, Richard A. Scheltema, Thijs J.G. Ettema, John van der Oost, David W. Taylor, Raymond H.J. Staals

## Abstract

Type III CRISPR-Cas systems provide a sequence-specific adaptive immune response that protects prokaryotic hosts against viruses and other foreign genetic invaders. These crRNA-guided Cas effector complexes bind and cleave complementary RNA targets. Specific target binding stimulates the Cas10 subunit to generate cyclic oligoadenylate (cOA) signaling molecules, that in turn allosterically activate proteins carrying cognate sensory domains: CARF or SAVED. Here, we characterize an elaborate set of genes associated with the type III-B CRISPR-Cas system from *Haliangium ochraceum*, which includes a signal transduction module of a CBASS defense system with two caspase-like proteases, SAVED-CHAT and PCaspase (Prokaryotic Caspase). We show that binding of a 3-nucleotide cOA (cA_3_) to the SAVED domain of SAVED-CHAT induces its oligomerization into long filaments that activate the proteolytic activity of the CHAT domain. Surprisingly, we find that activated SAVED-CHAT specifically cleaves and activates the second protease, PCaspase. In turn, activated PCaspase cleaves a multitude of other proteins, including a putative sigma factor and a PCaspase-inhibitor. We expressed the type III-B system and its associated genes in *E. coli* and observed a strong abortive phenotype when offering a complementary target RNA, but only in the presence of both SAVED-CHAT and PCaspase. Together, our findings show an intriguing cascade of proteolytic activities (conceptually similar to eukaryotic caspases) in this bacterial immune system that reveals yet another strategy to effectively defend against mobile genetic elements.

## Introduction

Type III CRISPR-Cas systems are adaptive immune systems in bacteria and archaea. These systems use CRISPR-derived RNA (crRNA) guides to target complementary nucleic acids of invading viruses and plasmids. Interestingly, type III systems have many unique features, including a rapidly expanding network of signal transduction pathways to trigger dormancy and cell death ^1–3^.

A typical type III operon encodes multiple Cas proteins that form a type III effector complex together with a mature crRNA guide. These complexes will bind complementary target RNA sequences, which initiate at an exposed seed region at the 3’ end of the crRNA guide. Seed binding initiates complete base pairing between the target RNA and the crRNA, resulting in the activation of Cas10, the characteristic multidomain subunit of the type III complex ^4^. The HD domain of activated Cas10 degrades ssDNA substrates in a non-sequence specific manner, whereas its Palm domain acts as a cyclase to convert ATP into signaling molecules called cyclic oligoadenylates (cOA), i.e. rings of 3-6 AMP moieties ^5–9^. These cOA signaling molecules activate a particular set of effector proteins carrying an appropriate cOA binding domain: CARF and SAVED proteins. These sensory domains are generally fused to a wide range of catalytic domains (e.g. RNases, DNases, NADases and toxins) ^10,11^. Over the last years, a handful of these CARF and SAVED proteins have been characterized, and despite their different activities, they all are geared towards killing the host, stopping the spread of the invading nucleic acid (e.g., phage progeny, plasmid propagation, etc.) in a process known as abortive infection. Recent work on type III systems indicated that proteases also play a role in type III immunity, as exemplified by the TPR-CHAT (Csx29) protease which associates with the type III-E complex, and a cOA-activated Lon-like protease (CalpL) in a type III-B system ^12,13^.

### A type III-B CRISPR-Cas/CBASS hybrid

We identified a set of genes that resides close to an operon encoding a type III-B CRISPR-Cas crRNA-guided protein complex (*Cmr1-6*) in the *Haliangium ochraceum* DSM 14365 genome (**Fig. 1a**). We observed a *SAVED-CHAT* gene, which encodes a fusion protein of a SAVED sensory domain and a CHAT domain (related to cysteine proteases that include the caspases, known to be involved in controlled cell death in eukaryotes). Further downstream, we observed a gene encoding a caspase-like cysteine protease (officially anotated as “Peptidase C14 caspase catalytic subunit p20”), which we named *PCaspase* (Prokaryotic Caspase). In between these genes, three genes are located that encode (i) a predicted sigma-factor, which we named *PCc-σ* (Prokaryotic Caspase-controlled sigma factor), (ii) a hypothetical gene, which we named *PCi* (Prokaryotic Caspase inhibitor, due to predicted structural homology with the CI-2 family of serine protease inhibitors), and (iii) a gene encoding a serine/threonine protein kinase, which we named *PCk (*Prokaryotic Caspase-controlled kinase). Notably, a similar operon appears to be encoded elsewhere on the chromosome of *H. ochraceum* (**Ext. Data Fig. 1a**).

**Fig. 1.**
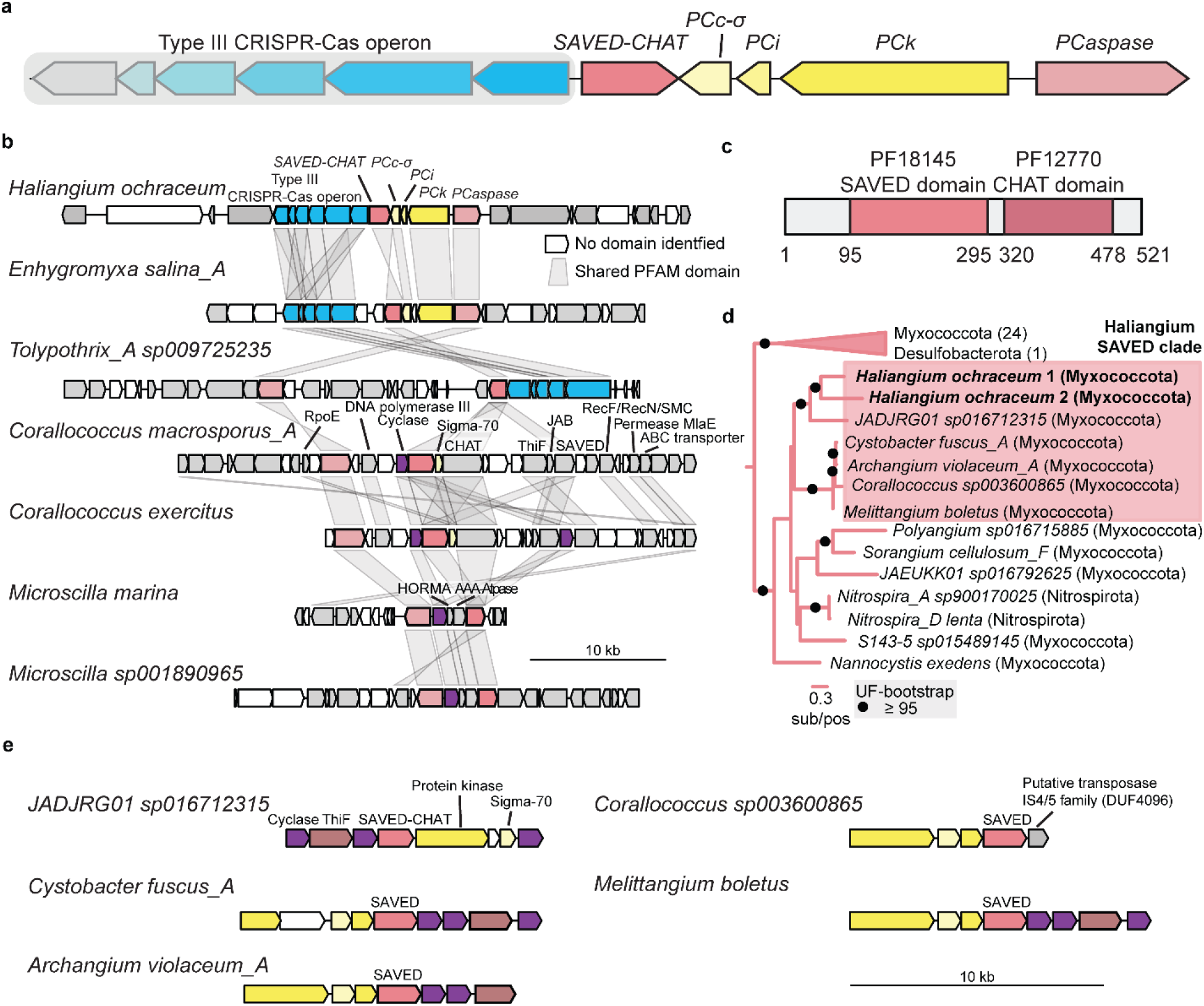
Type III CRISPR-Cas operon associated *SAVED-CHAT*/*PCaspase* gene cluster setup and CBASS origin. **a**, Schematic representation of the *H. ochraceum* type III-B CRISPR-Cas operon containing the Cmr1-6 genes encoding the crRNA-guided type III protein complex and its associated genes: *SAVED-CHAT, PCc-*σ*;, PCi, PCk* and *PCaspase (*locus tags Hoch_1313-1323). **b**, Genomic neighborhoods containing *SAVED-CHAT* and *PCaspase*. **c**, Domain architecture of SAVED-CHAT. **d**, SAVED domain phylogenetic midpoint-rooted tree. **e**, CBASS type operons containing *Haliangium* SAVED clade domains with associated proteins containing cyclase, protein kinase, Sigma-70 and transposase domains.

Next, we explored the co-occurrence of these genes in other prokaryotic genomes. We identified six additional bacterial genomes encoding *SAVED-CHAT* and *PCaspase* genes within the same gene cluster from the phyla Myxococcota (n=2), Bacteriodota (n=2) and Cyanobacteria (n=1) (**Fig. 1b**). Only the gene cluster of *Enhygromyxa salina*_*A* contained an identical gene set as *H. ochraceum*, with the exception of *PCi*, where homologs could not be identified in other prokaryotes. Type III CRISPR-Cas systems were present in the neighboring regions of two of the operons. In the remaining four operons CRISPR-Cas was missing in the neighboring genes, but genes encoding cGAS/DncV-like nucleotidyltransferase (CD-NTase) domains were found to cluster with *SAVED-CHAT*, indicative of CBASS defense systems.

Sensory domains like SAVED are often fused to different types of effector domains, implying a separate evolutionary origin of the SAVED and CHAT domains in *SAVED-CHAT* (**Fig. 1c**). We extracted the respective domains from representative prokaryotic and eukaryotic proteomes and calculated phylogenetic trees for each. The CHAT domains of the two *SAVED-CHAT* variants were only distantly related, indicating independent acquisition events (**Ext. Data Fig. 1b**). The *H. ochraceum* SAVED domains, however, formed a monophyletic clade (**Fig. 1d**) and were likely acquired once, after which a duplication resulted in two *SAVED* copies to which CHAT domains of different origin were added subsequently. This scenario is further supported by phylogenies of *PCaspase, PCc-σ*, and *PCk* (**Ext. Data Fig 1c-e**), which all support monophyly of the two respective copies. The SAVED domains are most closely related to those of other Myxococcota and might be part of a conserved system in these bacteria. We extracted the gene neighborhoods of *SAVED-CHAT* relatives and found that all but one contain a setup reminiscent of a CBASS defense system (**Fig. 1e**) composed of two to three putative cyclases, a protein kinase homolog of *PCk*, a *Sigma-70* like sigma factor and a ubiquitin activating *thiF* gene. Together, this suggests that cOA sensory and effector components of a CBASS system were co-opted in *H. ochraceum* to work in tandem with a type-III CRISPR-Cas system.

### SAVED-CHAT & PCaspase cleavage activity

We anticipated that SAVED-CHAT acts as a cOA-activated protease with a selective substrate repertoire similar to other members of CHAT cysteine proteases ^14^. To test this hypothesis, we purified all type III-associated proteins from *H. ochraceum* (except for PCk, which could not be cloned either individually or in combination with PCc-*σ* and PCi, likely due to toxicity) and conducted *in vitro* cleavage assays with SAVED-CHAT incubated with any of the three other proteins. The analysis indicates that SAVED-CHAT specifically cleaves PCaspase, in a cA_3_-dependent and co-factor independent manner (**Fig. 2a, Ext. Data Fig. 2, 3**), whereas no cleavage was observed for PCc-*σ* and PCi (**Fig. 2b**). SAVED-CHAT cleaved PCaspase into at least three defined fragments in addition to a myriad of products. A catalytically dead version of SAVED-CHAT (dSAVED-CHAT, H375A, C422A) showed no activity (**Ext. Data Fig. 2**). Lastly, we noticed that addition of cA_3_ prevented the migration of SAVED-CHAT in a native gel, indicating that SAVED-CHAT oligomerizes to form large complexes in the presence of cA_3_ (**Ext. Data Fig. 4**).

**Fig. 2.**
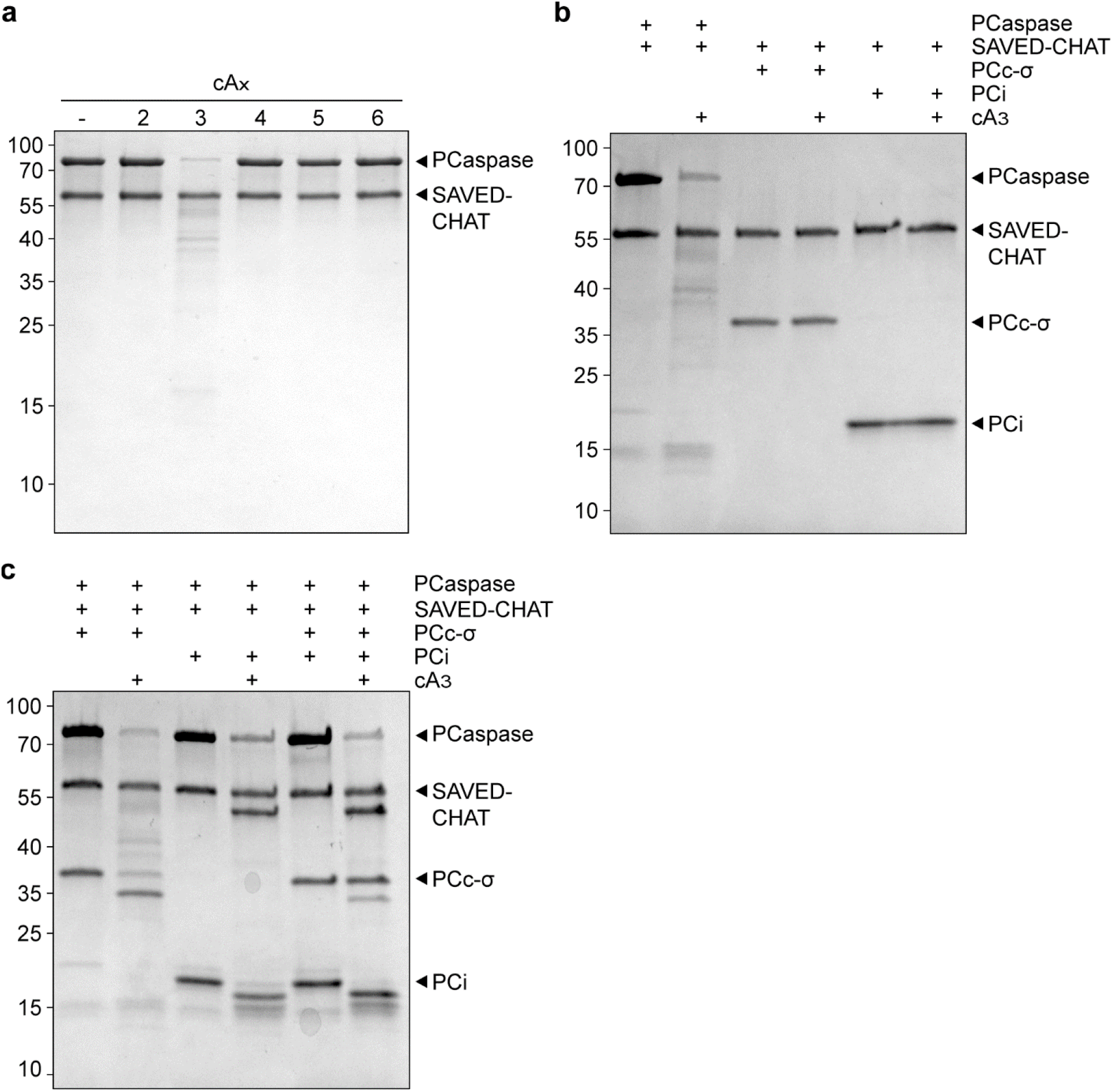
cA_3_-induced SAVED-CHAT and subsequent PCaspase activity. **a**, SDS-PAGE analysis of SAVED-CHAT cleavage, showing its dependency on cA_3_ for cleaving PCaspase. **b**, SDS-PAGE analysis of cleavage activity of activated SAVED-CHAT on PCc-*σ* and PCi. **c**, SDS-PAGE analysis of cleavage activity of activated PCaspase on PCc-*σ* and PCi. Activated PCaspase cleaves PCc-*σ* and PCi individually, PCc-*σ* is cleaved to a lesser extent when PCi is present.

Next, we performed protein cleavage assays with SAVED-CHAT and PCaspase in combination with either PCc-*σ* or PCi alone or with PCc-*σ* and PCi together (**Fig. 2c**). Cleavage of PCaspase by cA_3_-induced SAVED-CHAT resulted in the subsequent cleavage of PCc-*σ* and PCi into defined cleavage products. These results indicated that PCaspase cleavage by SAVED-CHAT activates its proteolytic activity. Interestingly, when PCi was present, one of the cleaved PCaspase fragments was stabilized and the smear of degradation products of PCaspase was absent, hinting at self-degradation. Furthermore, the efficiency of PCc-*σ* cleavage by PCaspase was decreased when PCi was included in the reaction, indicating that PCi acts as an inhibitor for PCaspase activity (**Fig. 2c**). Indeed, similar PCaspase-mediated PCc-*σ* cleavage reactions with increasing PCi concentrations showed increasing rates of inhibition (**Ext. Data Fig. 5**).

We hypothesized that PCaspase might have a broader substrate specificity than the associated genes alone. We provided a biologically unrelated casein protein as substrate to test this idea. Indeed, activated PCaspase leads to the degradation of casein in a co-factor independent manner, while SAVED-CHAT by itself does not (**Fig. 3a, Ext. Data Fig. 6**). Furthermore, a catalytic mutant of PCaspase (dPCaspase, H79A, C146A) shows no activity (**Fig. 3a**). A PCi titration experiment demonstrates that PCi inhibits PCaspase activity on casein, showing that this inhibitory effect is independent from PCc-*σ* (**Ext. Data Fig. 7**).

**Fig. 3.**
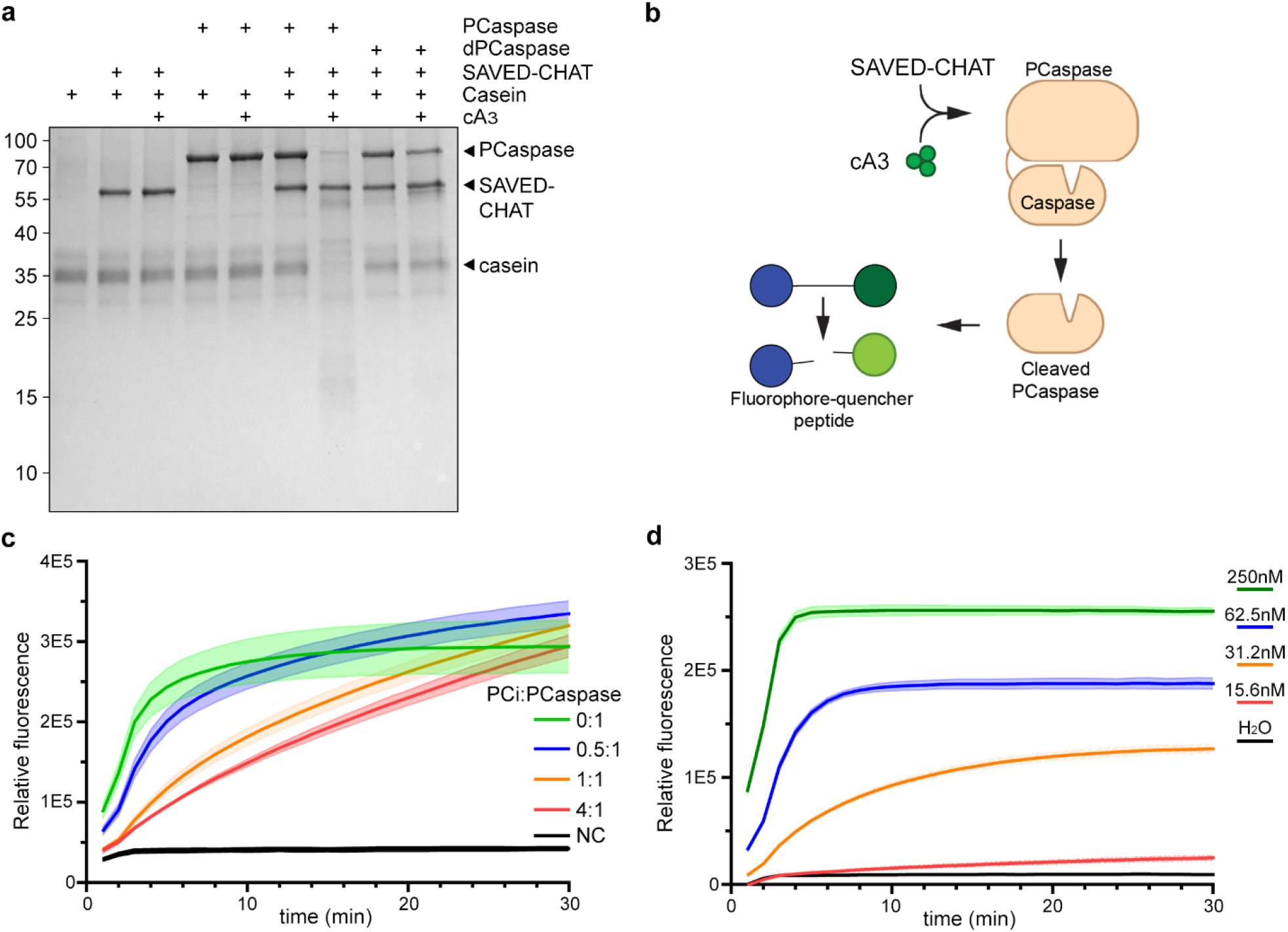
PCaspase activity on non-associated substrates. **a**, SDS-PAGE analysis of the proteolytic activation of PCaspase by SAVED-CHAT, resulting in degradation of the casein substrate, in contrast to activated SAVED-CHAT by itself or the dPCaspase catalytic mutant (H79A, C146A). **b**, Schematic overview of PCaspase activity assay using a fluorescent peptide reporter. c, Increasing the molar ratio (PCi:PCaspase) of PCi reduces the cleavage activity of a FAM-peptide substrate by activated PCaspase. The faded colors represent the standard error of the mean (n=3). d, Sensitivity for cA_3_ of the FAM-peptide visualization method is 15.6 nM. The faded colors represent the standard error of the mean (technical replicates, n=3).

To monitor PCaspase activity in real-time, a small fluorophore-quencher peptide was provided as a substrate, which is specifically cleaved by activated PCaspase after the cA_3_ induced cascade is initiated (**Fig. 3b, Ext. Data Fig. 8**). With this, the inhibitory effect of PCi on PCaspase can be monitored in real-time, which shows a clear, inverse relationship between reaction kinetics and amount of PCi (**Fig. 3c**). To probe the sensitivity and kinetics of this detection method to cA_3_, a concentration range was used to determine near instant signal generation in the presence of >31.2 nM and a limit of detection of 15.6 nM (**Fig. 3d**).

### SAVED-CHAT and PCaspase activation leads to a strong abortive infection phenotype

Due to the apparent broad-substrate specificity of PCaspase, we hypothesized that the sequential activation of *H. ochraceum* type III CRISPR-Cas, SAVED-CHAT, and eventually, PCaspase by a target RNA would lead to an abortive infection phenotype. For this, we constructed the pHochTypeIII plasmid, which expresses *H. ochraceum cmr1-6* (forming the type III complex), *csb2* (a *cas6* homologue for processing of the pre-crRNA), and a minimal CRISPR array containing two repeats and a spacer (**Ext. Data Fig. 9a**)^15^. The SAVED-CHAT and PCaspase genes were expressed in different combinations from the pEffector plasmid (**Ext. Data Fig. 9b**). NucC (a cA_3_-responsive nuclease, known to cause abortive infection) from *E. coli* and an empty vector were used as positive and negative controls, respectively (**Ext. Data Fig. 9b**)^16^. Target and non-target plasmids were constructed, carrying a protospacer sequence complementary to the spacer, expression of which was induced by IPTG (**Ext. Data Fig. 9c**)^17^.

Target or non-target plasmid was transformed into *E. coli* BL21-AI harbouring pHochTypeIII and one of various pEffector plasmids **(Fig. 4a)**. Subsequent 10-fold serial dilutions were plated on glucose-or IPTG-containing plates to repress or induce, respectively, the target or non-target RNA, and CFUs were scored to calculate transformation efficiencies. First, we demonstrate that the type III system produces cA_3_ upon target RNA recognition and leads to an abortive infection phenotype in the presence of NucC (**Fig. 4b**)^16^. Furthermore, only in the condition where both SAVED-CHAT and PCaspase are present, and upon the expression of the target RNA, transformation efficiency is reduced by six orders of magnitude (**Fig. 4b**). With this, we confirm that type III from *H. ochraceum* produces (at least) cA_3_ and is able to mount a very strong abortive infection phenotype in combination with its effector proteins SAVED-CHAT and PCaspase.

**Fig 4.**
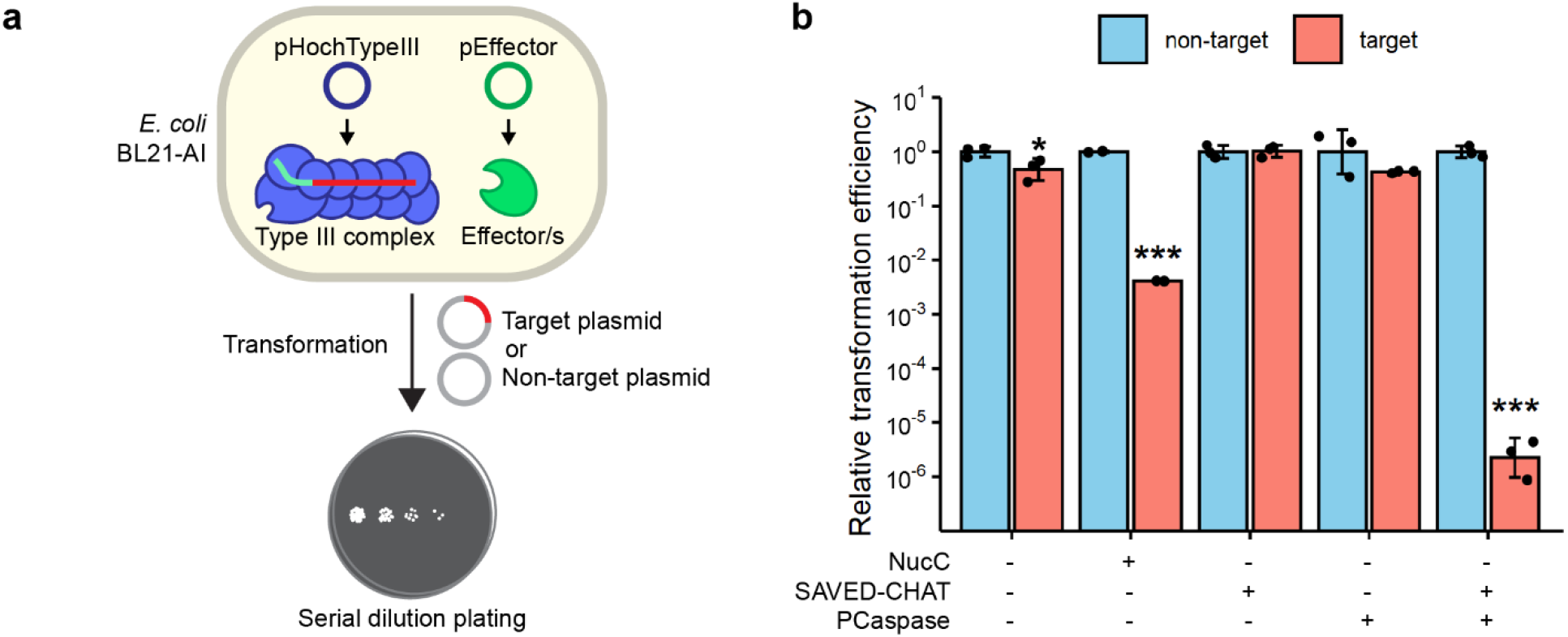
PCaspase activation severely reduced the transformation efficiency of a target plasmid. **a**, Schematic overview of the experimental setup in *E. coli* BL21-AI. **b**, Transformation efficiencies of target or non-target plasmid in *E. coli* co-expressing *H. ochraceum* type III CRISPR-Cas complex and different combinations of effector proteins as indicated. Statistical significance was calculated using one-sided unpaired Welch’s t-test. * - *p* < 0.05, ** - *p* < 0.005, *** - *p* < 0.0005 (n=3).

### Structural basis for SAVED-CHAT activation

To understand the structural basis for cA_3_-induced activation, we determined a cryo-EM structure of the activated SAVED-CHAT in complex to cA_3_. Size-exclusion chromatography (SEC, data not shown) indicated that while SAVED-CHAT is monomeric in solution, whereas cA_3_ binding induces formation of large, polydisperse oligomers. This phenomenon was also observed with native gel electrophoresis (**Ext. Data Fig. 3**). Prolonged incubation of cA_3_ with SAVED-CHAT resulted in the formation of large, polymorphous aggregates that were not amenable to structural determination. However, by vitrifying immediately after mixing SAVED-CHAT and cA_3_ we were able to obtain a structure of the filament at a global resolution of 3.1 Å.

SAVED-CHAT oligomerization results in long, curved filaments with the SAVED domain on the inside and the CHAT domains at the periphery, with a curvature of ∼10° between each monomer (**Fig. 5a**). SAVED-CHAT monomers assemble via head-to-tail oligomerization, with a single cA_3_ bound at the interface between two SAVED domains, forming ‘singlet’ filaments. We additionally observed partial ‘doublets’ where two antiparallel filaments make a cross-fiber interaction spanning up to three SAVED-CHAT monomers (**Fig. 5a**).

**Fig. 5.**
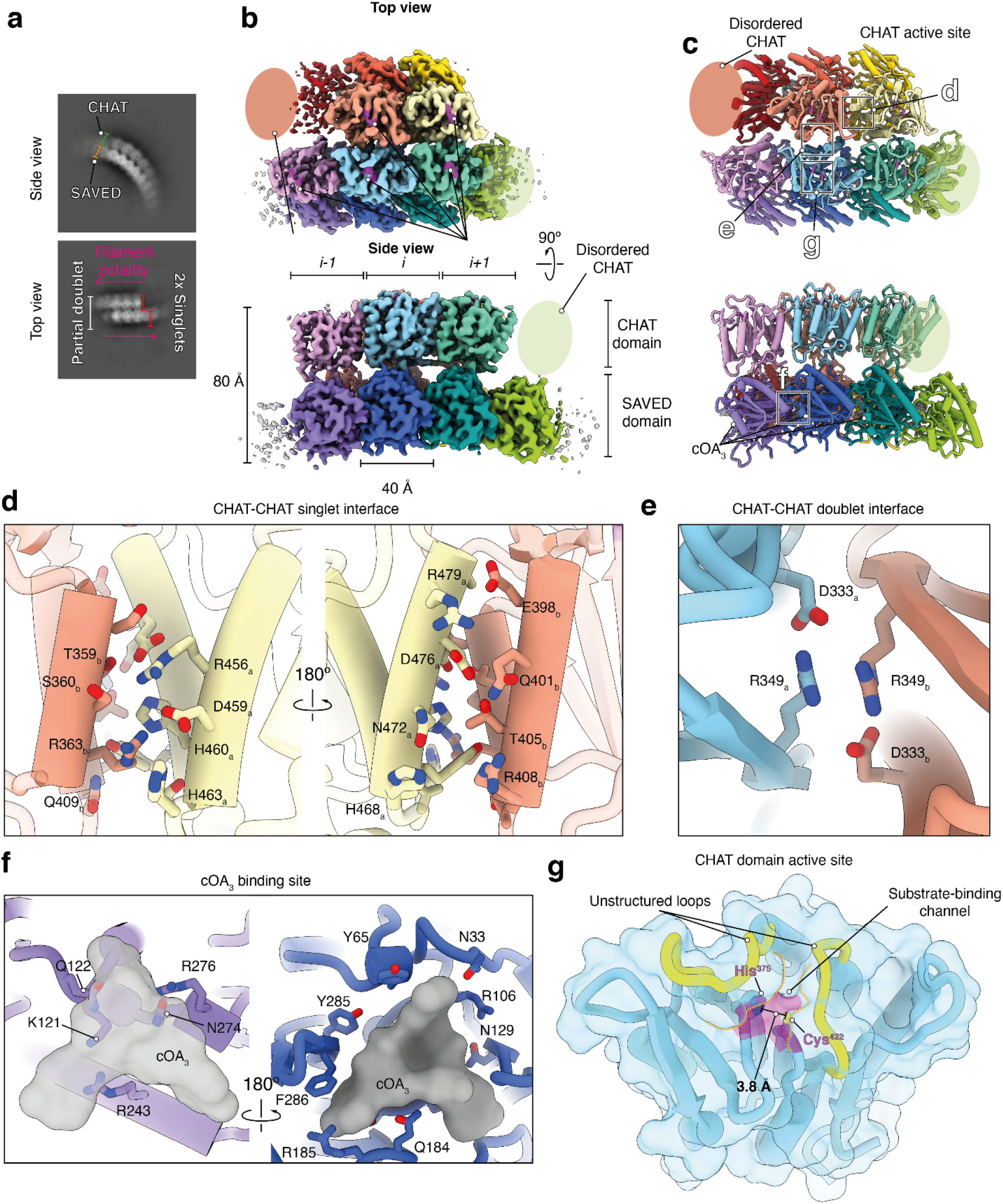
Structural basis for SAVED-CHAT activation by cA_3_. **a**, 2D class averages of SAVED-CHAT bound to cA_3_, showing side and top views. In the side view, the filament curvature is evident, with the CHAT domain at the periphery of the arch. In the top view, singlet filaments form a partial inter-filament doublet, with two singlets running in opposite polarities forming cross-fiber contacts spanning ∼3 monomers. **b & c**, 3.1 Å-resolution cryo-EM reconstruction and model of the cA_3_-bound SAVED-CHAT filament. In some monomers, the CHAT domain is disordered and absent from the reconstruction (red and light green monomers). **d**, Close-up view of the CHAT-CHAT singlet intra-filament interface, consisting of a four-helix bundle. **e**, Close-up view of the CHAT-CHAT doublet inter-filament interface. **f**, cA_3_ binding site, at the interface between adjacent SAVED domains. **g**, CHAT domain active site. The residues H375 and C422 that comprise catalytic dyad (magenta) are 3.8 Å apart and located within the substrate-binding channel beneath two unstructured gating loops (yellow).

In the ‘singlet’ regions of our map, the CHAT domains were poorly resolved and often absent due to molecular motion (**Fig. 5b, c**). Analysis of AlphaFold2 models of monomeric SAVED-CHAT revealed a high degree of flexibility between the two globular domains, with an unstructured linker (residues 297-304) acting as a hinge to accommodate a maximum displacement of up to 35 Å (**Ext. Data Fig. 10**). It is likely that the strain induced by the filament curvature prevents more than three successive CHAT domains from forming stable inter-fiber contacts.

The intra-filament head-to-tail CHAT-CHAT interface is comprised of a four-helix bundle with two helices from each protomer (**Fig. 5d**). This bundle is held together through a network of predominantly electrostatic interactions. The CHAT-CHAT inter-filament doublet interface is mediated by an unusual π-π stacking interaction between two R349 residues (that is, the same residue from different monomers), reinforced by additional electrostatic contacts (**Fig. 5e**).

The cA_3_ is buried within the intra-filament interface between two SAVED domains, and participates in a plethora of hydrogen bonds, electrostatic and stacking interactions (**Fig. 5f**). This network of contacts suggests that cA_3_ acts as a molecular glue to bridge SAVED domain intra-filament interactions, which subsequently provides a platform for CHAT domain rigidification, doublet formation and ultimately substrate capture. This is a distinct mechanism of activation from the TIR-SAVED filaments which assemble a composite active site across two adjacent TIR domains within a filament which assembles upon cA_3_-binding, or the CRISPR-associated Lon protease (CalpL) activation that appears to have a concentration-dependent effect, reflecting the mechanistic diversity of cA-responsive ancillary factors 13,18.

Within the resolved doublet CHAT domains, the active site is positioned underneath two unstructured loops which create a narrow substrate-binding channel (**Fig. 5g**). This channel is wide enough to accommodate unstructured peptides, but would likely be inaccessible to highly structured peptides. The Cys422-His375 catalytic dyad are aligned and positioned ∼4Å apart, confirming that the cA_3_-bound filament represents the active conformation of SAVED-CHAT. Comparison with the CHAT domain of the activated type III-E protein TPR-CHAT bound to proteolytic substrate Csx30, revealed a near-identical active site configuration where the catalytic dyad is aligned, and the substrate-binding channel is exposed (**Ext. Data Fig. 10**).

## Discussion

Caspases are a family of cysteine proteases, that generally use a catalytic cysteine residue to target proteins by cutting a peptide bond. Caspases of animals generally cut after an aspartate, whereas caspase variants of fungi and plants (meta-caspases) cut after arginine or lysine ^19–24^. In eukaryotes, these caspases are key players in signal transduction pathways that often trigger programmed cell death (apoptosis) ^22,25^. These apoptotic pathways consist of two classes of caspases: initiators and executioners, the former ones proteolytically activating the latter ones to trigger further downstream events ^26^. Our work highlights a conceptual similarity on this theme in prokaryotes: a cOA-sensory SAVED domain that activates a cysteine protease (caspase-like CHAT, analogous to an initiator), which in turn activates a second cysteine protease (PCaspase, analogous to an executioner) that acts as a central node to set in motion a cascade of events. PCaspase appears to modify downstream targets that are predicted to trigger dormancy and/or cell death.

The model in **Fig. 6** summarizes our current understanding of the CRISPR-Cas type III-B system in *H. ochraceum* and its effector components, where detection of target RNA results in the generation of cA_3_ (not shown in schematic) ^8,9^. The synthesized cA_3_ binds to the sensory domain of SAVED-CHAT. Acting as a molecular glue, cA_3_ participates in a multitude of interactions that initially stabilize SAVED:SAVED dimerization (**Fig. 4**). The subsequent multimerization of the unusual antiparallel SAVED-CHAT doublet filament results in the activation of the CHAT domains, an unprecedented mechanism for CARF and SAVED effector proteins (**Fig. 4**). The activated CHAT domains in turn cleave PCaspase, which then becomes an active protease, cleaving at least PCc-*σ* and PCi into defined products (**Fig. 2**). In contrast to the specific cleavage of PCaspase by SAVED-CHAT, PCaspase has a broader substrate repertoire, as reflected by the cleavage of the casein and FAM-peptide substrates *in vitro* (**Fig. 3)**. This is further supported by the strong abortive infection phenotype observed when PCaspase is activated *in vivo* (**Fig. 4**), where proteolytic activity of other host factors most likely leads to the significantly reduced transformation efficiency.

**Fig. 6.**
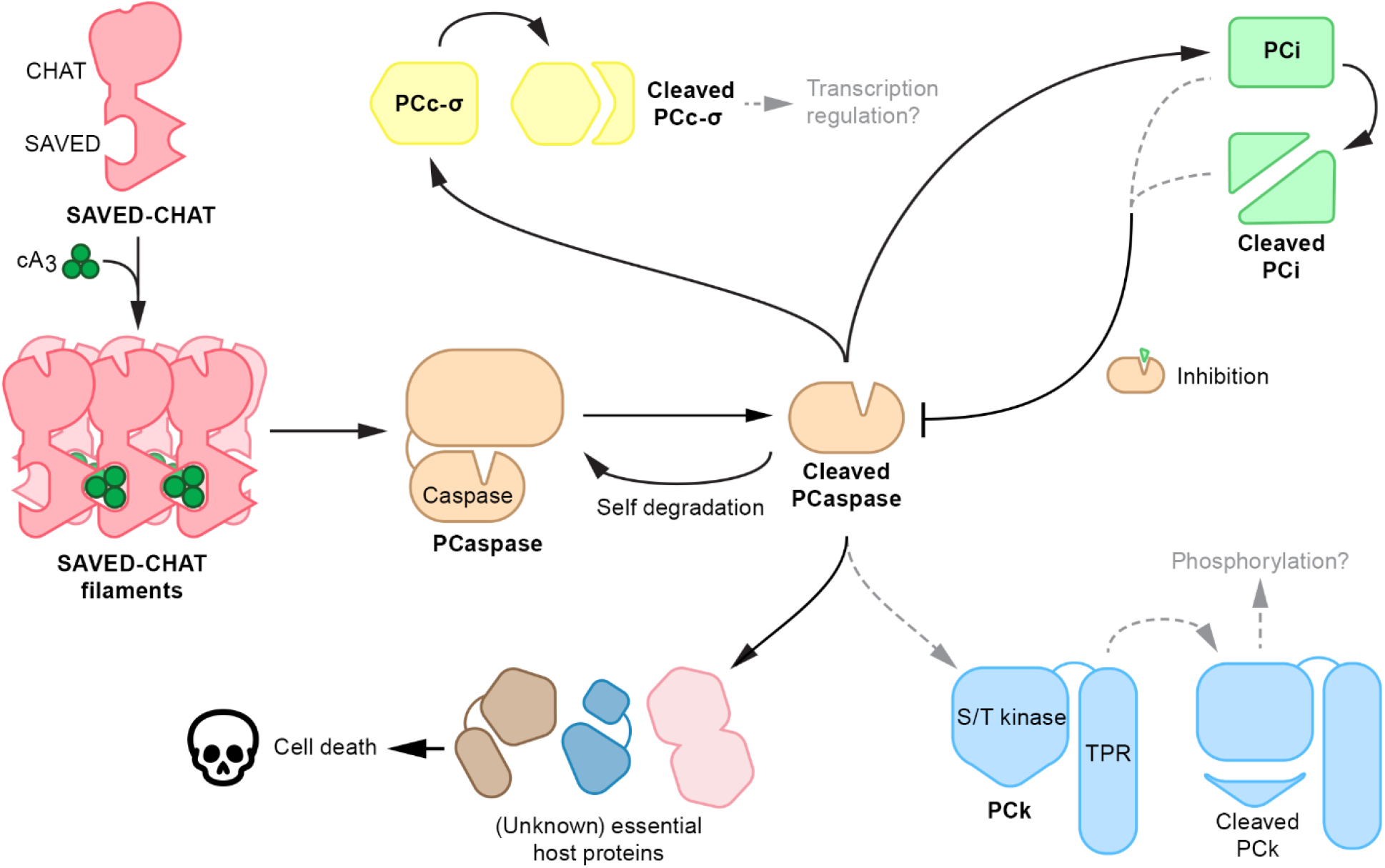
Model of the type III-B CRISPR-Cas system and its associated genes in *H. ochraceum*. Binding of cA_3_ to the SAVED domain of SAVED-CHAT induces oligomerization that activates its CHAT domain. Activated SAVED-CHAT cleaves PCaspase, which subsequently mediates further downstream events by cleaving PCi (inhibitor of PCaspase), PCc-*σ* (transcriptional response?) and potentially also PCk (phosphorylation / toxicity?) and/or other essential host proteins. Hypothesized events are in grey.

Interestingly, we found that PCi is a potent inhibitor of PCaspase on all tested substrates *in vitro*, hinting at an auto-regulated feedback loop to specifically control PCaspase activity (**Fig. 2**). However, we currently cannot verify whether the pre-or post-cleaved PCi fulfils this role (**Fig. 6**). Furthermore, the predicted sigma factor PCc-*σ* might be involved in coordinating additional defence strategies through transcriptional regulation, as previously suggested for CASP-*σ* in type III-E ^27^. In the latter study it was shown that CASP-*σ* is part of a sub-complex together with Csx30 (a hypothesized toxin) and Csx31, both lacking any similarity to either PCi or PCk. Upon target RNA binding by the type III-E protein complex, the associated protease TPR-CHAT cleaves Csx30 and releases CASP-*σ* for transcriptional regulation of target genes that may include Cas1, hinting at a connection between interference and spacer adaptation in type III CRISPR-Cas systems. Other recent work on the type III-B system from *Sulfurihydrogenibium sp*. showed that CalpL (a Lon-like protease fused to a SAVED domain) forms a complex together with CalpT (a potential anti-sigma factor) and CalpS (a sigma factor) ^13^. Upon cOA signalling, CalpL oligomerizes and subsequently cleaves CalpT, releasing a subcomplex consisting of a CalpT fragment and CalpS that is thought to regulate gene expression by RNAP.

Which target genes PCc-*σ* from our type III-B system will regulate is an interesting question for future studies to address. It is apparent, however, that the type III-B system studied here involves a different set of genes, including two different proteases (SAVED-CHAT and PCaspase), a predicted transcriptional regulator (PCc-*σ*), an auto-inhibitor (PCi) and a potential toxic protein kinase (PCk).

Probably due to toxicity issues, all attempts to (co)-purify PCk or include it in the *in* vivo experiments have thus far not been successful. As shown in **Fig. 6**, we hypothesize that activated PCaspase might proteolytically modify and activate PCk, setting yet another signal transduction pathway in motion. Bacterial Ser/Thr kinases have been demonstrated to act in stress response and anti-phage defense by phosphorylating several target proteins, at least in some cases hampering growth ^28,29^. Hence, it might be that the anticipated cytotoxicity of PCk reflects its Ser/Thr kinase activity, which may in turn play a role in instigating abortive infection, either directly in response to a PCaspase-mediated cleavage, or indirectly through induction of its transcription (PCc-*σ*). There is also still more to understand about how the activity of PCaspase is suppressed by PCi. It is tempting to speculate that this apparent feedback mechanism may allow for down-regulating the stress response at the level of a reversible dormancy state, rather than going irreversibly to cell death.

In recent years, an unprecedented diversity has been elucidated of type III, cOA-based signal transduction pathways ^1,2^. In all variants that have recently been described, many relevant details remain to be discovered. Most notably, the full impact of further downstream events following activation of CRISPR-associated proteases is not fully understood and is very likely to vary between different CRISPR-Cas systems. The variant described in this work is an intriguing example of evolutionary tinkering of prokaryotic immune systems that has resulted in a set of caspase-like proteases to convert a stress signal to trigger cell death.

## Acknowledgements

R.H.J.S. is supported by a VIDI grant (VI.Vidi.203.074) from the Dutch Research Council (NWO), J.v.d.O. by an ERC Advanced Grant (ERC-AdG-834279) from the European Research Council, and a Spinoza grant from the Dutch Research Council (SPI 93-537). T.J.G.E. received funding from the European Research Council (CoG grant 817834), the Dutch Research Council (VICI grant VI.C.192.016) and the Volkswagen Foundation (grant 96725). D.W.T.is supported in part by the National Institute of General Medical Sciences of the National Institutes of Health (R35GM138348), the Welch Foundation Research Grant (F-1938), and a Robert J. Kleberg, Jr. and Helen C. Kleberg Foundation Medical Research Grant. D.W.T is an American Cancer Society Research Scholar supported by the American Cancer Society (RSG-21-050-01-DMC). We thank Rob Joosten for technical support on some of the experiments.

## Author contributions

J.A.S., J.v.d.O. and R.H.J.S conceived of and designed the study. S.K. and T.J.G.E. performed gene cluster and phylogenetic analyses. J.A.S, J.P.K.B, C.R.P.S, C.Y, A.M.A, S.H.P.P, C.P, An.B, A.B and R.A.S performed experiments and analyses. J.A.S, J.P.K.B, J.v.d.O, K.S, T.J.G.E, D.W.T and R.H.J.S wrote the manuscript with significant input from other authors.

## Data statement

The data that support the findings of this study are available from the corresponding author upon reasonable request.

## Methods

### Computational identification of SAVED-CHAT/PCaspase gene clusters

We downloaded predicted protein coding sequences for all species representatives of archaea and bacteria from the genome taxonomy database (GTDB) (PMID: 30148503) release 207 (https://data.gtdb.ecogenomic.org/releases/release207/207.0/genomic_files_reps/gtdb_proteins_aa_reps_r207.tar.gz). According to annotated Pfam (PMID: 33125078) domains in PCaspase (locus tag Hoch_1323) and SAVED-CHAT (locus tag Hoch_1319), we searched with Pfam hidden Markov models (HMM) of Caspase (PF00656), SAVED (PF18145) and CHAT (PF12770) domains against the species representative proteomes with the “hmmsearch” program in HMMER v3.1b2 (PMID: 22039361) with default parameters. The identified candidate proteins were reannotated using InterProScan v5.50-84 (PMID: 24451626) with Pfam domains (-appl Pfam) to remove false positive hits. We screened 62,291 bacterial and 3,412 archaeal species representative genomes for domain fusions of SAVED and CHAT, and a caspase domain within 10 genes distance. For each of the three Pfam domains, we defined candidate gene clusters. Final gene clusters were formed by merging overlapping candidate gene clusters and annotated with Pfam domains. Gene clusters were visualized with GenoPlotR v0.8.11 (PMID: 20624783).

### Phylogenetic analysis of the SAVED-CHAT/PCaspase gene cluster proteins

For *SAVED-CHAT* and *PCaspase*, we performed domain specific phylogenetic analysis. We extracted all regions with these domain annotations from the GTDB prokaryotic species representative genome dataset. We additionally performed Pfam domain annotation as described above for the eukaryotic protein sequences in the eggNOG v5 (PMID: 30418610) database (http://eggnog5.embl.de/download/eggnog_5.0/e5.proteomes.faa) and extracted the respective protein domains. We then merged prokaryotic and eukaryotic protein domains and performed multiple sequence alignment (MSA) with FAMSA v1.6.2 (PMID: 27670777). After gentle trimming with trimAl (PMID: 19505945) v1.4.rev15 (-gt 0.02), we calculated approximate maximum likelihood (ML) trees with FastTree v2.1.11 (PMID: 20224823). We extracted focal clades for respective *H. ochraceum* proteins in the two gene clusters. The respective focal clade sequences were aligned with MAFFT L-INS-i (PMID: 23329690) v7.427 and we inferred ML trees with IQ-TREE (PMID: 32011700) v2.0.3 (-m MFP –bb 1000). For PC*c-σ* and PCk (locus tags: Hoch_1320 and Hoch_1322, respectively) we searched the GTDB representative species protein dataset with MMseqs2 (PMID: 29035372) v14.7e284 (--max-seqs 2000) to identify the highest scoring homologs. We aligned sequences with FAMSA, trimmed MSAs with trimAL (-gt 0.02) and calculated trees with FastTree. Trees were visualized with FigTree v1.4.4 (http://tree.bio.ed.ac.uk/software/figtree/).

### Plasmids and strains for protein purification

Electrocompetent *E. coli* DH5α were used for general cloning and plasmid propagation. Full-length genes of SAVED-CHAT, PCaspase, PCc-*σ* and PCi from *H. ochraceum* DSM 14365 (NCBI reference sequence NC_013440.1) were codon-harmonized for expression in *E. coli* BL21 (DE3) and synthesized with an N-terminal Strep-tag II by either Integrated DNA Technologies (IDT) or Twist Bioscience. The synthetic genes were then cloned into the expression vector pJS-BCD under an IPTG-inducible T7 promoter. Catalytically dead mutants of SAVED-CHAT (dSAVED-CHAT) and PCaspase (dPCaspase) were generated by PCR mutagenesis. All expression plasmids (**Ext. Data Table 1**) were transformed into electrocompetent *E. coli* BL21 (DE3) for protein expression and purification.

### Protein purification

For recombinant expression and purification of SAVED-CHAT, PCaspase (and mutants hereof), PCc-*σ* and PCi, the expression vector pJS-BCD was used (**Ext. Data Table 1**). *E*.*coli* BL21 (DE3) cells transformed with the expression vector were inoculated in 1.5 L lysogeny broth (LB) and grown at 37°C with 150 rpm shaking. After an optical density (O.D. 600 nm) of 0.6-0.7 was reached, a cold shock on ice for 30-60 min was performed. Protein expression was induced by adding 0.5 mM IPTG (isopropyl β-D-1-thiogalactopyranoside). Subsequently, cells were grown at 16 °C for 18-24 hours and harvested via centrifugation. Pelleted cells were resuspended in 20-30 mL Wash Buffer (150 mM NaCl, 100 mM Tris, pH 8.0) and a cOmplete protease inhibitor tablet (Roche) was added. Cells were lysed by sonication and lysate was clarified by centrifugation and subsequent filtration (0.45uM). The supernatant was applied to a StrepTrap XT 5 mL column (Cytiva) and the bound protein was eluted with Elution Buffer (Wash Buffer + 50 mM biotin). Finally, a Superdex 200 (Cytiva) size exclusion chromatography column was used with Wash Buffer as eluate. After confirmation with SDS-PAGE analysis, protein fractions were pooled and concentrated to a working stock of 5 μM.

### Protein cleavage activity assay

All cleavage assays were performed in a buffer containing 125 mM NaCl, 10 mM Tris-HCl (pH 8.0), and 1 mM DTT. For SAVED-CHAT, PCaspase (and mutants thereof), PCc-*σ*, and PCi, a final protein concentration of 0.5 μM was added, unless otherwise specified. Depending on the assay, a final concentration of 1 μM cA_3_ (Biolog.de), 5 mM EDTA, 5 mM EGTA, 2 mM MgCl2, 2 mM CaCl, 1 mM ATP and 0.1 mg/mL casein was added. After incubation for 1h at 35°C, SDS loading dye was added and incubated for a further 5 min at 95°C. SDS-PAGE analysis using Coomassie blue staining was utilized to visualize the results (Biorad Gel Doc XR).

### SAVED-CHAT complex formation on native PAGE

For native PAGE analysis of SAVED-CHAT complex formation, 1 μM SAVED-CHAT was incubated with or without 1 μM cA_3_ for 1h at 35°C in a buffer containing 125 mM NaCl, 10 mM Tris-HCl (pH 8.0), and 1 mM DTT. Afterwards, the reaction was run on a native 4-20% polyacrylamide gel, stained with Coomassie blue, and visualized (Biorad Gel Doc XR).

### Real-time PCaspase activity assay

SAVED-CHAT/PCaspase activity assays were conducted *in vitro* in activity buffer (125 mM NaCl, 10 mM Tris, 1 mM DTT, pH 8.0) to which different combinations of component were added: SAVED-CHAT (0.5 μM), PCaspase (0.5 μM), cA_3_ (15.6 nM to 1 μM), and/or FAM-peptide substrate (5 μM) (Eurogentec AS-60579-01). Assays were incubated for one hour at 37C with a FAM channel measurement at 1 min intervals in a Thermo Scientific Quantstudio 1 RT-qPCR instrument running Quantstudio Design & Analysis software (v1.5.2). Data was visualized using Graphpad Prism 9 (n=3) with standard error depiction.

### Plasmid interference assay

Electrocompetent *E. coli* BL21-AI carrying pHochTypeIII and one of the pEffectors (**Ext. Data Fig. 9a**,**b**) were prepared for target/non-target plasmid transformation (**Ext. data Table 1**). For pHochTypeIII, *H. orchaceum* type III-B *cmr1-6, csb2*, and a minimal CRISPR array carrying one spacer sequence were placed individually under the control of T7 promoters. For the pEffectors, the various effector genes were cloned into one operon and placed under the control of a constitutively expressing lacUV5 promoter. For the target/non-target plasmids, a non-coding RNA was placed under the control of a trc promoter and lacO (**Ext. Data Fig. 9c**)^17^. This non-coding RNA carries the protospacer targeted by the spacer of the crRNA guide encoded by the CRISPR array on pHochTypeIII.

The transformations were carried out in triplicate, using 100 ng of target or non-target plasmid, by electroporation (BTX electroporation system). After transformation, all cells were recovered in 1mL of LB at 37°C for one hour. Ten-fold dilutions were plated on LB-agar medium containing 0.2% arabinose, 34 μg/mL chloramphenicol, 50 μg/mL carbenicillin, 50 μg/mL kanamycin, and either 1 mM IPTG or 0.2% glucose to induce or repress target RNA transcription, respectively. Finally, the plates were incubated overnight at 30°C, and transformation efficiencies were quantified. Data was analyzed and visualized using R version 4.3.0, with statistical significance calculated by one-sided unpaired Welch’s t-test.

### Structural analyses

10 µM SAVED-CHAT was mixed with 125 µM cA_3_. 2.5 µl of complex was immediately applied to C-flat grids (1.2/1.3, 300 mesh) which had been plasma-cleaned for 30 seconds in a Solarus 950 plasma cleaner (Gatan) with a 4:1 ratio of O_2_/H_2_. Grids were blotted with Vitrobot Mark IV (Thermo Fisher) for 6 seconds, blot force 0 at 4°C & 100% humidity, and plunge-frozen in liquid ethane. Data were collected on a FEI Glacios cryo-TEM equipped with a Falcon 4 detector. Data was collected in SerialEM, with a pixel size of 0.94 Å, a defocus range of -1.5 --2.5 µm, and a total exposure time of 15s resulting in a total accumulated dose of 40 e/Å^2^ which was split into 60 EER fractions. Motion correction, CTF estimation and particle picking was performed on-the-fly using cryoSPARC Live v4.0.0-privatebeta. Data were collected on a FEI Glacios cryo-TEM equipped with a Falcon 4 detector, as described for the binary complex. In total, 2,550 movies were collected. All subsequent data processing was performed in cryoSPARC v3.2. 1,229,738 particle co-ordinates were picked, selected and subjected to 2D classification. Multiple rounds of *ab-initio* modelling and heterogeneous refinement resulted in a subset of 39,001 particles, which yielded a 3.1 Å-resolution which was used for modelling. For modelling, multiple AlphaFold2 models of the individual SAVED and CHAT domains were rigid-body fitted into the map. cA_3_ was built in Coot, and other adjustments were performed in Isolde, before running real-space refinement in Phenix. All structural figures and movies were generated using ChimeraX.

## References

1. van Beljouw, S. P. B., Sanders, J., Rodríguez-Molina, A. & Brouns, S. J. J. RNA-targeting CRISPR–Cas systems. Nat Rev Microbiol 21, (2022).

2. Steens, J. A., Salazar, C. R. P. & Staals, R. H. J. The diverse arsenal of type III CRISPR-Cas-associated CARF and SAVED effectors. Biochem Soc Trans 50, 1353–1364 (2022).

3. Steens, J. A., van der Oost, J. & Staals, R. H. J. Compact but mighty: Biology and applications of type III-E CRISPR-Cas systems. Mol Cell 82, 4405–4406 (2022).

4. Steens, J. A. et al. SCOPE enables type III CRISPR-Cas diagnostics using flexible targeting and stringent CARF ribonuclease activation. Nat Commun 12, 1–12 (2021).

5. Elmore, J. R. et al. Bipartite recognition of target RNAs activates DNA cleavage by the Type III-B CRISPR–Cas system. Genes Dev 30, 447–459 (2016).

6. Estrella, M. A., Kuo, F. T. & Bailey, S. RNA-activated DNA cleavage by the Type III-B CRISPR–Cas effector complex. Genes Dev (2016) doi:10.1101/gad.273722.115.

7. Kazlauskiene, M., Tamulaitis, G., Kostiuk, G., Venclovas, C. & Siksnys, V. Spatiotemporal Control of Type III-A CRISPR-Cas Immunity: Coupling DNA Degradation with the Target RNA Recognition. Mol Cell 62, 295–306 (2016).

8. Niewoehner, O. et al. Type III CRISPR-Cas systems produce cyclic oligoadenylate second messengers. Nature 548, 543–548 (2017).

9. Kazlauskiene, M., Kostiuk, G., Venclovas, C., Tamulaitis, G. & Siksnys, V. A cyclic oligonucleotide signaling pathway in type III CRISPR-Cas systems. Science (1979) 357, 605–609 (2017).

10. Makarova, K. S., Anantharaman, V., Grishin, N. V., Koonin, E. V. & Aravind, L. CARF and WYL domains: Ligand-binding regulators of prokaryotic defense systems. Front Genet 5, 1–9 (2014).

11. Makarova, K. S. et al. Evolutionary and functional classification of the CARF domain superfamily, key sensors in prokaryotic antivirus defense. Nucleic Acids Res 48, 8828–8847 (2020).

12. van Beljouw, S. P. B. et al. The gRAMP CRISPR-Cas effector is an RNA endonuclease complexed with a caspase-like peptidase. Science (1979) 373, 1349–1353 (2021).

13. Christophe Rouillon, Niels Schneberger, Haotian Chi, Katja Blumenstock, Stefano Da Vela, Katrin Ackermann, Jonas Moecking, Martin F. Peter, Wolfgang Boenigk, Reinhard Seifert, Bela E. Bode, Jonathan L. Schmid-Burgk, Dmitri Svergun, Matthias Geyer, M. F. W. & G. H. Antiviral signaling by a cyclic nucleotide activated CRISPR protease. Nature 1–23 (2022) doi:10.1038/s41586-022-05571-7.

14. Aravind, L. & Koonin, E. V. Classification of the caspase-hemoglobinase fold: Detection of new families and implications for the origin of the eukaryotic separins. Proteins: Structure, Function and Genetics 46, 355–367 (2002).

15. Shangguan, Q., Graham, S., Sundaramoorthy, R. & White, M. F. Structure and mechanism of the type I-G CRISPR effector. Nucleic Acids Res 50, 11214–11228 (2022).

16. Lau, R. K. et al. Structure and Mechanism of a Cyclic Trinucleotide-Activated Bacterial Endonuclease Mediating Bacteriophage Immunity. Mol Cell 77, 723–733.e6 (2020).

17. Ichikawa, H. T. et al. Programmable type III-A CRISPR-Cas DNA targeting modules. PLoS One 12, 1–12 (2017).

18. Hogrel, G. et al. Cyclic nucleotide-induced helical structure activates a TIR immune effector. Nature 608, 808–812 (2022).

19. Alnemri, E. S. et al. Human ICE/CED-3 protease nomenclature. Cell 87, 171 (1996).

20. Talanian, R. V. et al. Substrate specificities of caspase family proteases. Journal of Biological Chemistry 272, 9677–9682 (1997).

21. Carmona-Gutierrez, D., Fröhlich, K. U., Kroemer, G. & Madeo, F. Editorial: Metacaspases are caspases. Doubt no more. Cell Death Differ 17, 377–378 (2010).

22. Cohen, G. M. Caspases: Executioners of Apoptosis. Pathobiology of Human Disease: A Dynamic Encyclopedia of Disease Mechanisms 16, 145–152 (2014).

23. Watanabe, N. & Lam, E. Two Arabidopsis metacaspases AtMCP1b and AtMCP2b are arginine/lysine-specific cysteine proteases and activate apoptosis-like cell death in yeast. Journal of Biological Chemistry 280, 14691–14699 (2005).

24. Vercammen, D. et al. Type II metacaspases Atmc4 and Atmc9 of Arabidopsis thaliana cleave substrates after arginine and lysine. Journal of Biological Chemistry 279, 45329–45336 (2004).

25. Fan, T. J., Han, L. H., Cong, R. S. & Liang, J. Caspase family proteases and apoptosis. Acta Biochim Biophys Sin (Shanghai) 37, 719–727 (2005).

26. Boatright, K. M. & Salvesen, G. S. Mechanisms of caspase activation. Curr Opin Cell Biol 15, 725–731 (2003).

27. Strecker, J., Demircioglu, F. E., Li, D., Faure, G. & Max, E. RNA-activated protein cleavage with a CRISPR-associated endopeptidase. Science (1979) 7450, 1–16 (2022).

28. Gratani, F. L. et al. E. coli Toxin YjjJ (HipH) Is a Ser / Thr Protein Kinase That Impacts Cell Division, Carbon Metabolism, and Ribosome Assembly. mSystems (2022).

29. Depardieu, F. et al. A Eukaryotic-like Serine/Threonine Kinase Protects Staphylococci against Phages. Cell Host Microbe 20, 471–481 (2016).

30. Fang, C. et al. Structures and mechanism of transcription initiation by bacterial ECF factors. Nucleic Acids Res 47, 7094–7104 (2019).

