## Extended Data Figures for "Type III-B CRISPR-Cas signaling-based cascade of proteolytic cleavages"

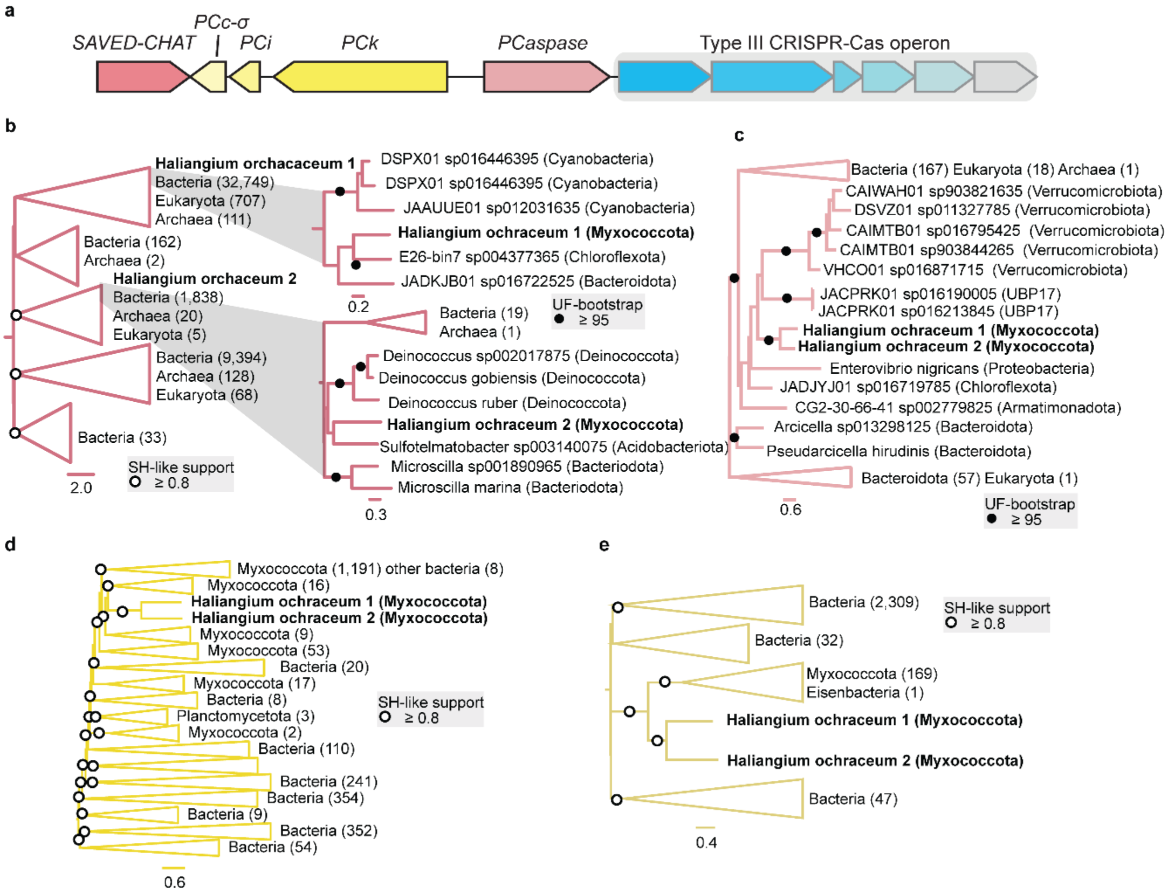


**Ext. Data Fig. 1. Second SAVED-CHAT/PCaspase like gene cluster and protein phylogenies. a,** Schematic representation of the second *H. ochraceum* type III CRISPR-Cas operon and its associated genes: *SAVED-CHAT*, *PCc-σ*, *PCi*, *PCk*, and *PCaspase* (locus tags Hoch_5578-5588). **b,** CHAT domain general and individual focal clade phylogenetic trees. **c,** Phylogenetic tree of caspase domain focal clade. **d and e,** Phylogenetic trees of *PCk* and *PCc-σ*, respectively. All trees are midpoint-rooted, scale bars indicate substitutions per position in the alignment.

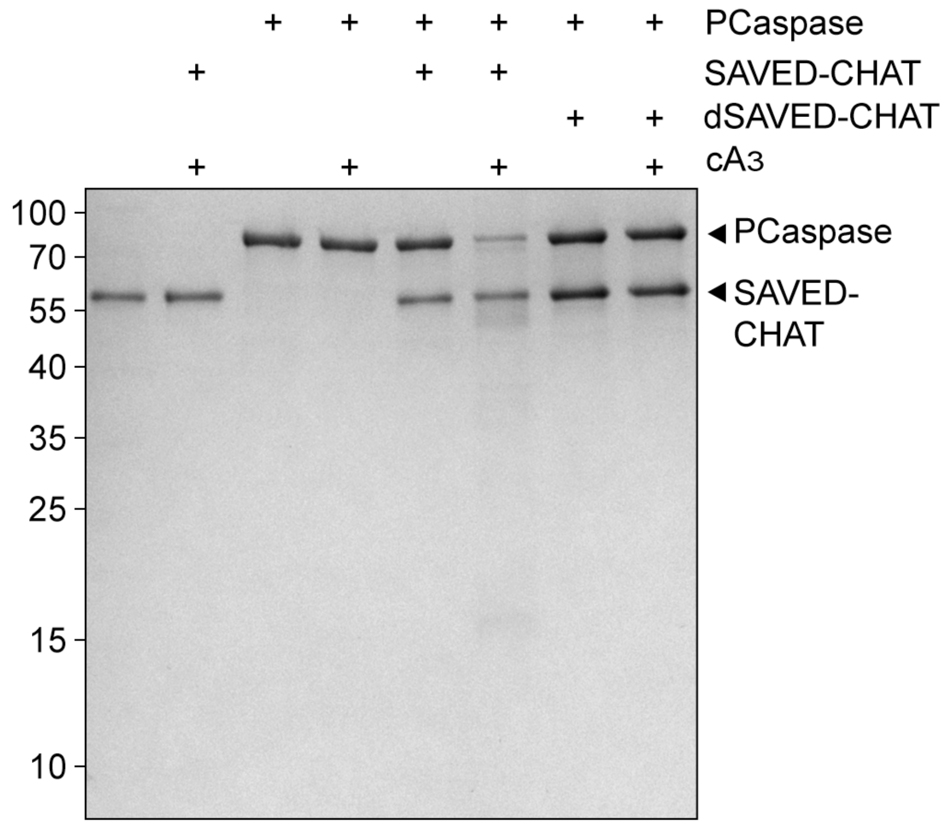


**Ext. Data Fig. 2. SDS-PAGE analysis of (d)SAVED-CHAT activity.** PCaspase is specifically cleaved by $cA_3$-activated SAVED-CHAT, opposed to the SAVED-CHAT H375A/C422A catalytic mutant.

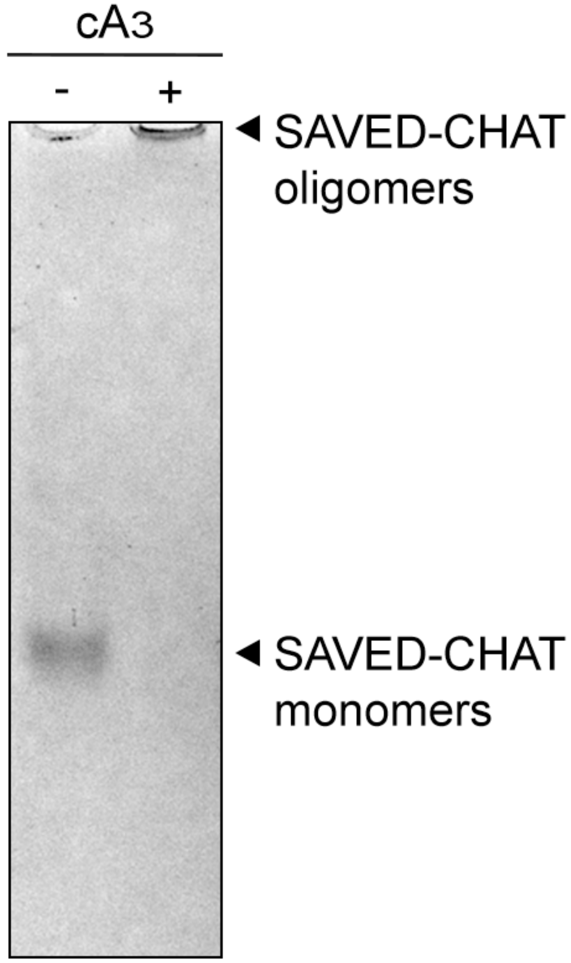


**Ext. Data Fig. 3. Native PAGE analysis of SAVED-CHAT after incubation with cA3.** Oligomerization of SAVED-CHAT monomers hampers migration into a native PAGE gel in a cA3-dependent maner.

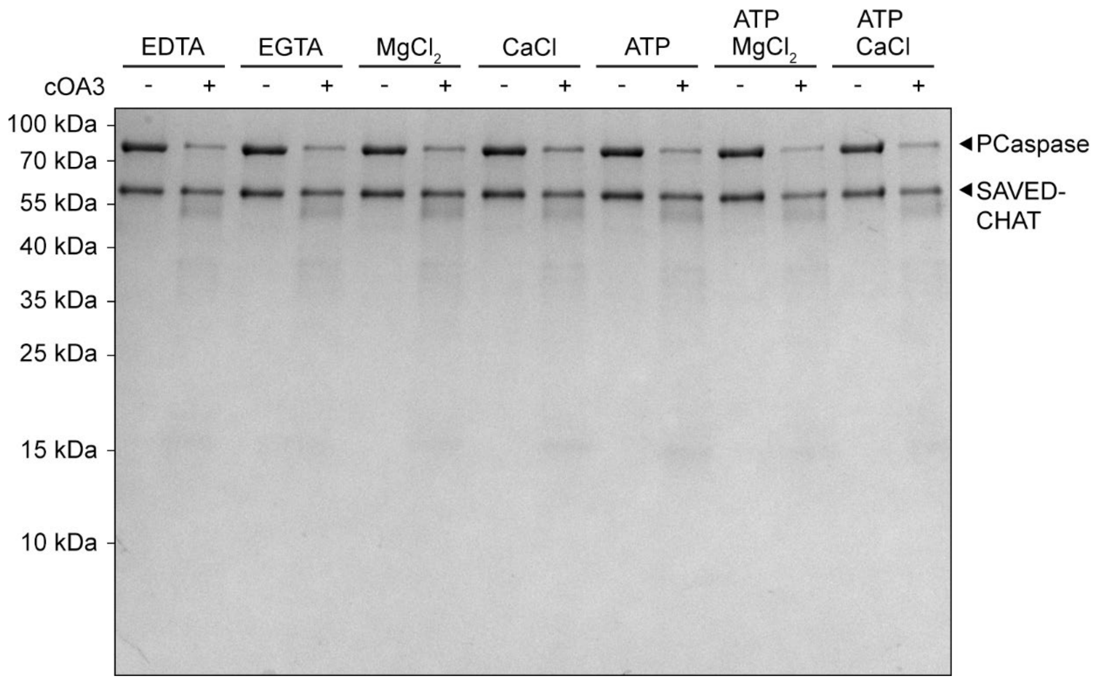


**Ext. Data Fig. 4 SDS-PAGE analysis of co-factor requirements of PCaspase cleavage by SAVED-CHAT.** There is no apparent dependence on a co-factor or the catalytic activity of SAVED-CHAT on PCaspase when activated by $cA_3$.

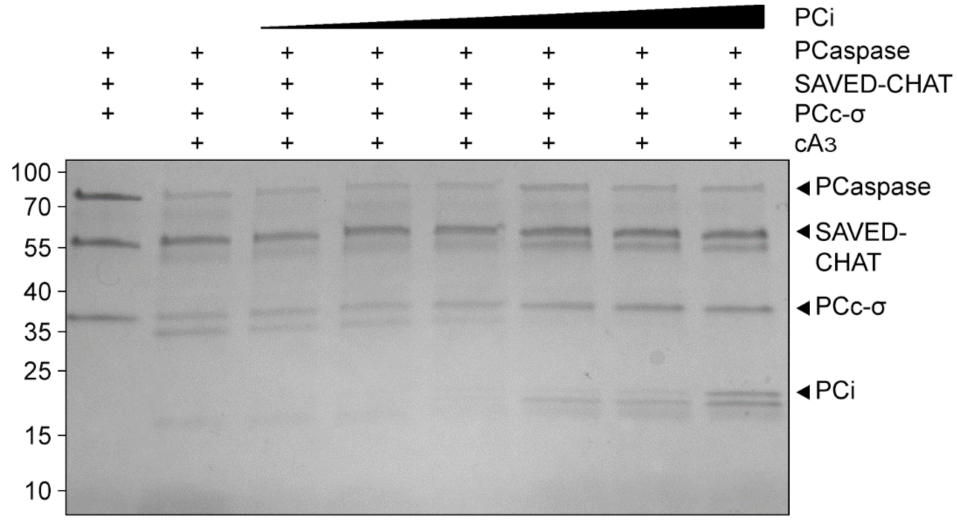


**Ext. Data Fig. 5. SDS-PAGE analysis of the inhibitory effects of PCi on PCaspase cleavage activity on PCc-σ**. An increasing concentration of PCi reduces the ability PCaspase to cleave PCc-σ, cleavage of PCi is also uncomplete. Molar ratios of PCi:PCaspase range from 1:20 – 2:1.

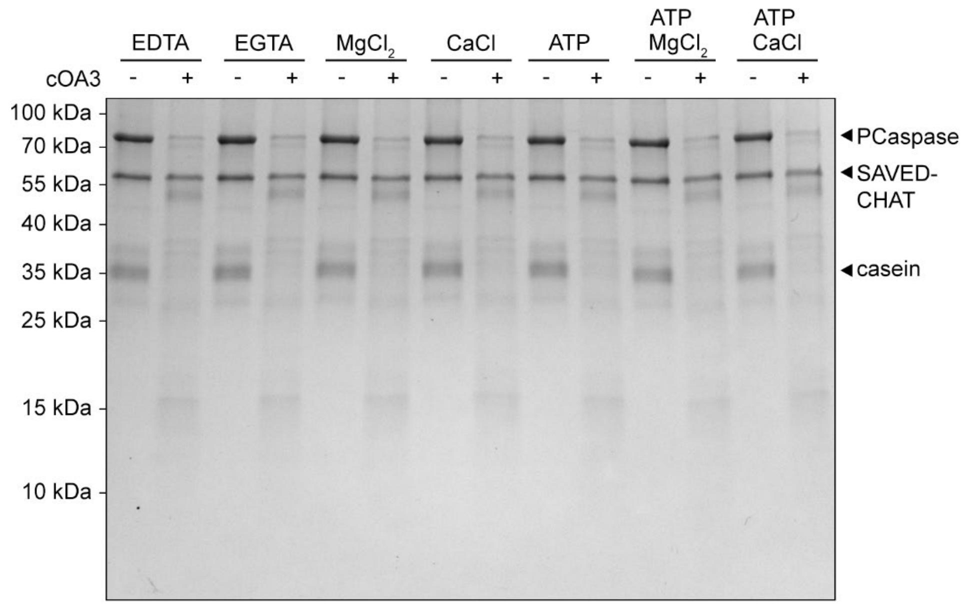


**Ext. Data Fig. 6. SDS-PAGE analysis of co-factor requirements ofPCaspase cleavage activity of casein.** SDS-PAGE protein cleavage assays with SAVED-CHAT, PCaspase and casein, demonstrating that the cleavage activity on casein of activated PCaspase does not require co-factors.

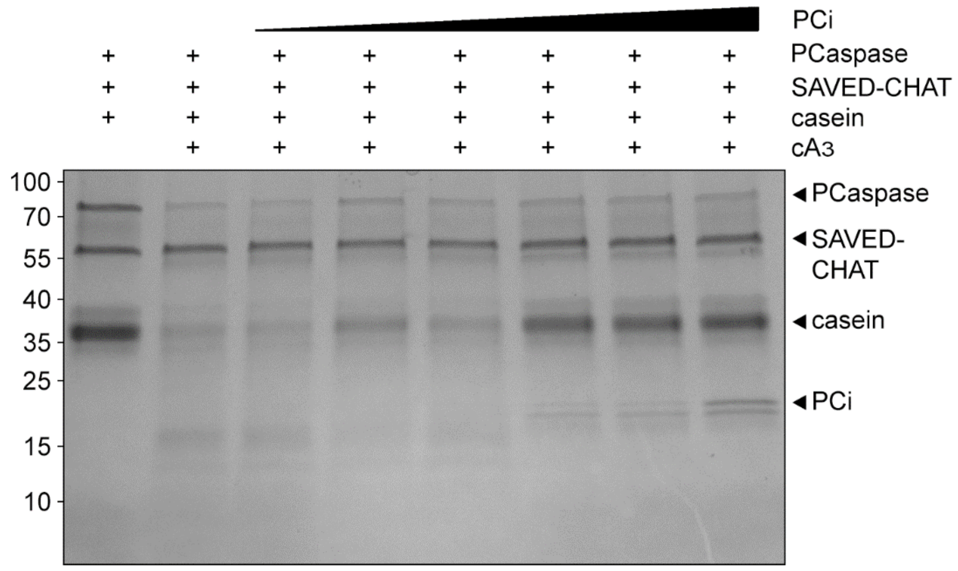


**Ext. Data Fig. 7. SDS-PAGE analysis of the inhibitory effects of PCi on casein cleavage by activated PCaspase.** An increasing concentration of PCi reduces the ability of activated PCaspase to cleave casein and complete cleavage of PCi. Molar ratios of PCi:PCaspase range from 1:20 – 2:1.

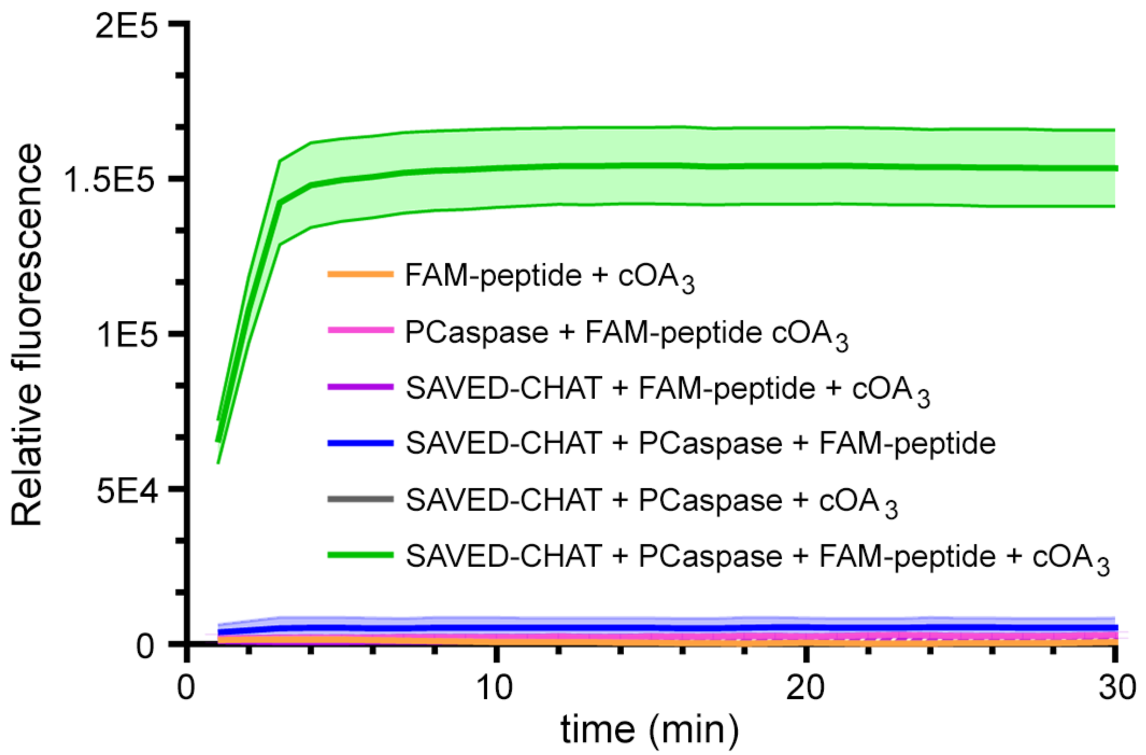


**Ext. Data Fig. 8. FAM-peptide is specifically cleaved by activated PCaspase.** Various combinations of proteins in graph are represented by colored lines. The transparent bands represent the standard error of the mean (technical replicates, n=3).

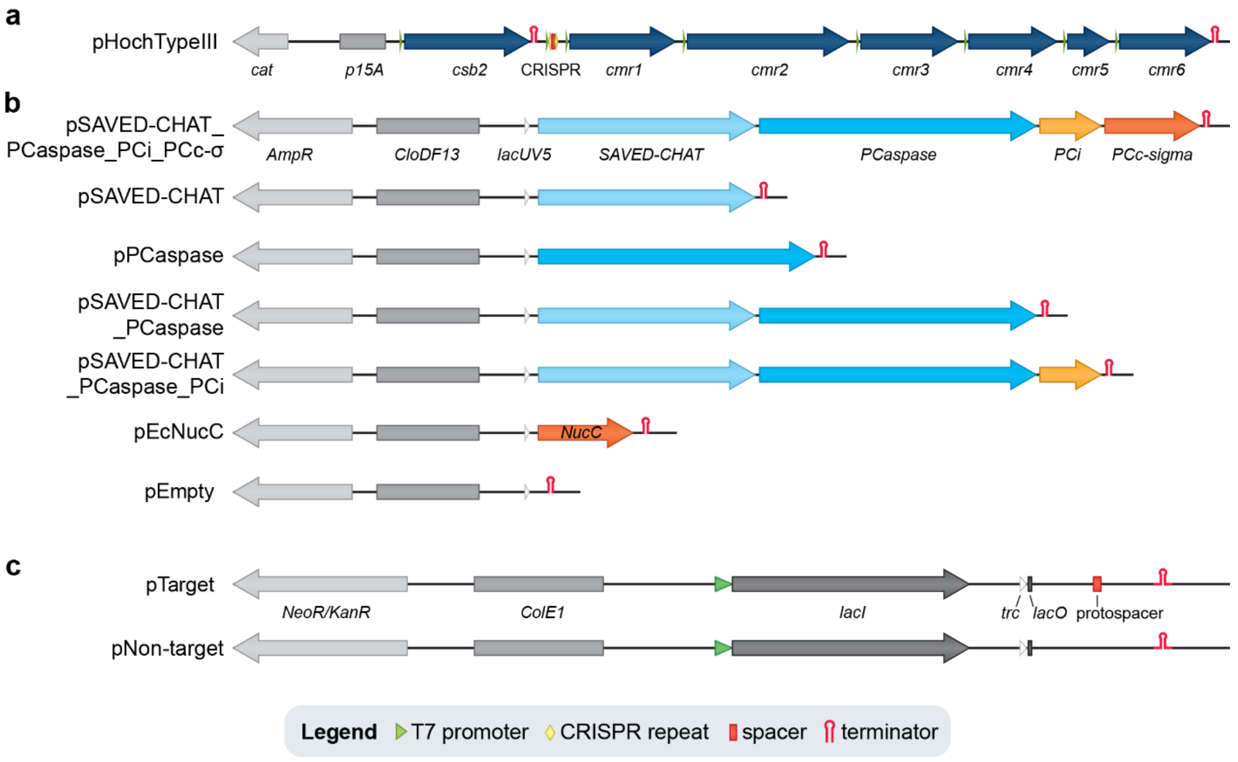

**Ext. Data Fig. 9. Linearized maps of the plasmids used in the plasmid challenge assay. a,** Map of the *H. ochraceum* type III CRISPR-Cas expression plasmid pHochTypeIII, with *cmr1*-*cmr6* from the *SAVED-CHAT* genomic neighborhood, *csb2* from a co-occurring type I-G system, and the associated CRISPR array with a single spacer sequence targeting a protospacer on pTarget. **b,** Maps of the effector expression plasmids, showing the different combinations of effectors used in the study. pEcNucC and pEmpty were used as positive and negative controls. **c,** Maps of the pTarget and pNon-target, having identical backbones except for a protospacer on pTarget.

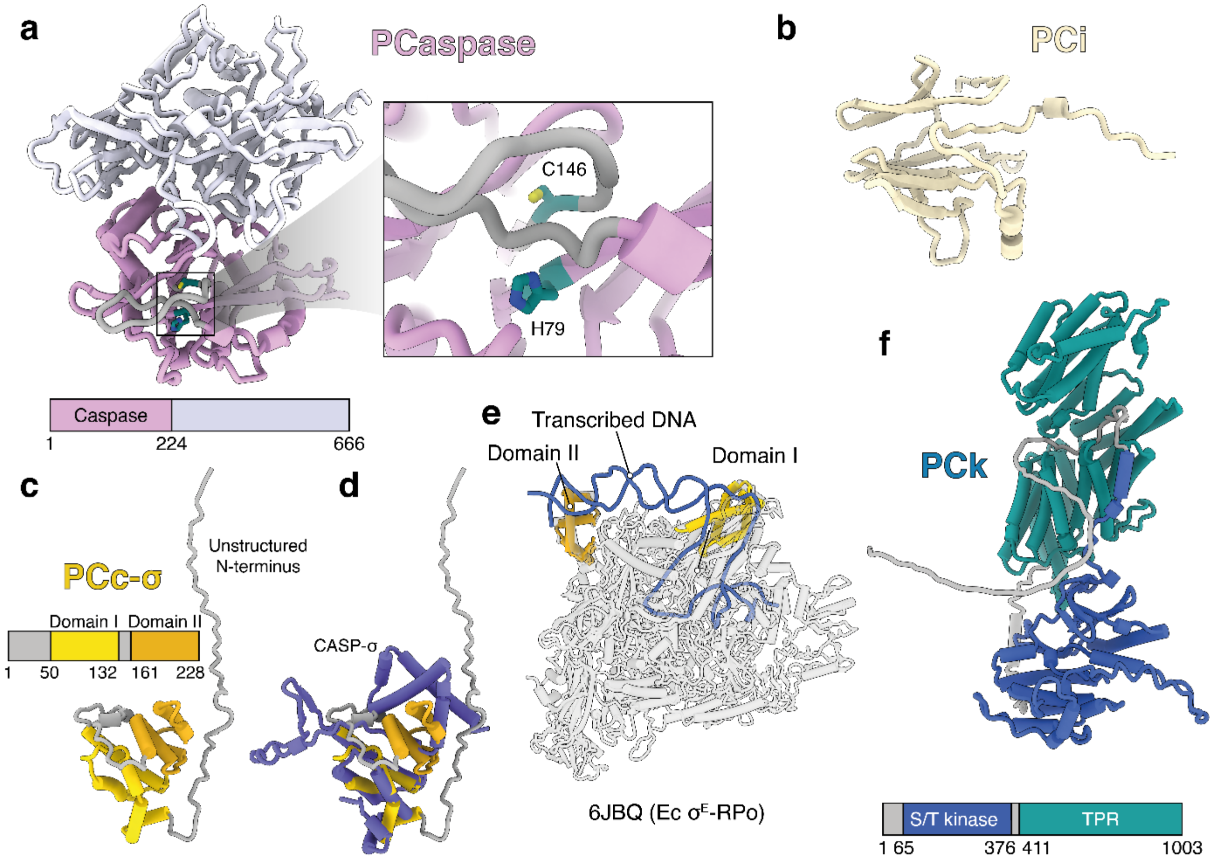


**Ext. Data Fig. 10 AlphaFold2 analysis of other factors encoded within the type III-associated genes a,** Predicted model of PCaspase. The proteolytic CHAT domain is shown in pink, and the C-terminal putative regulatory domain is shown in light grey. Right: close-up view of catalytic dyad H79 and C146 in an autoinhibited conformation, with a peptide occluding the active site. **b**, Precited model of PCi. **c**, Predicted model of PCc-σ. Two sigma factor domains (Domain I and Domain II) are shown in yellow and brown. These domains are attached by a flexible linker. **d**, alignment of PCc-σ with CASP-σ [27]. **e**, Alignment of two PCc-σ domains with *E. coli* RNA polymerase bound to $\sigma^E$ (PDB ID 6JBQ) [30]. **f**, Predicted model of PCk. S/T kinase domain is shown in blue, and TPR is in dark cyan.

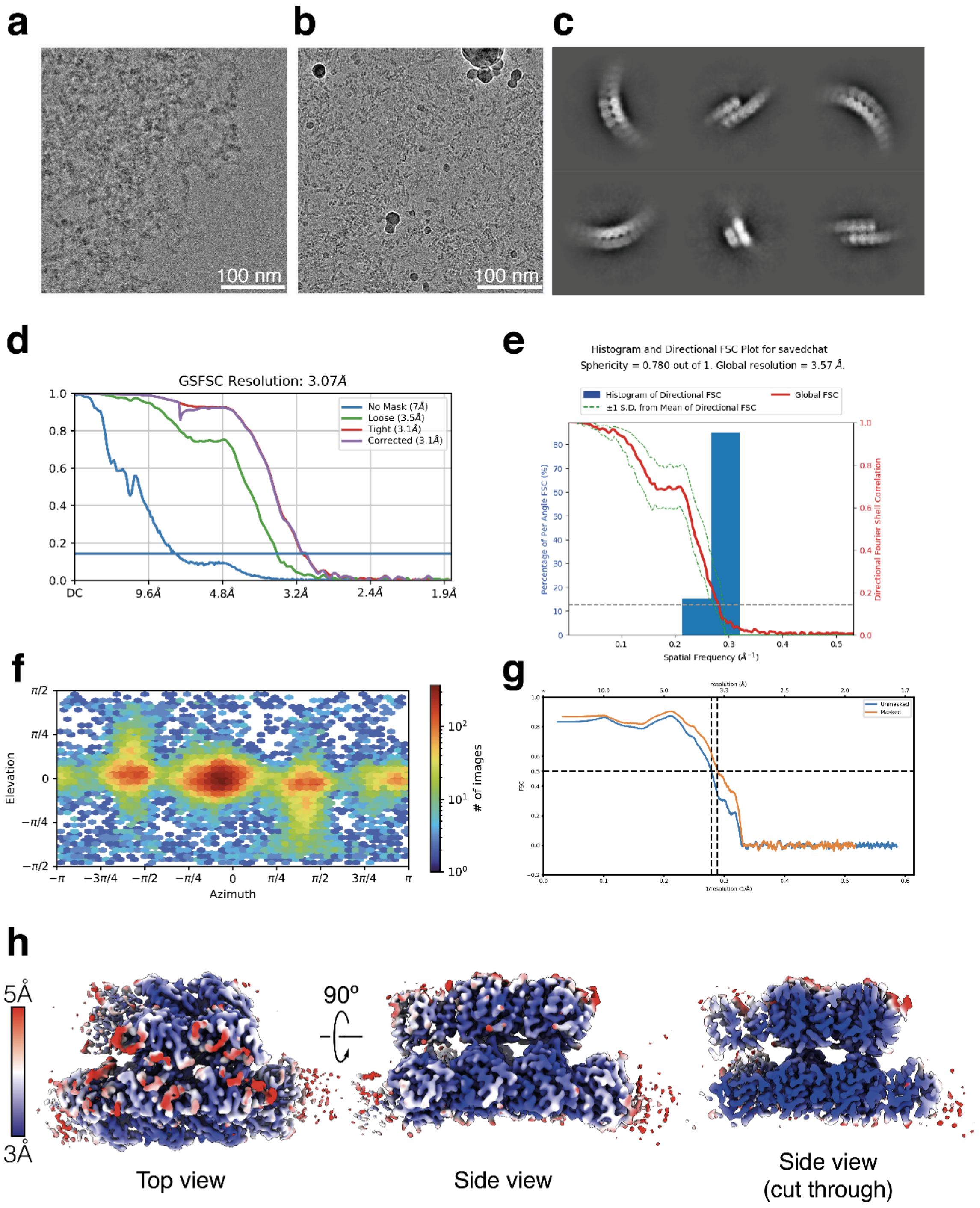


**Ext. Data Fig. 11. Cryo-EM data collection and validation. a,** Cryo-EM micrograph of SAVED-CHAT incubated for 1 hour at 25ºC with $cA_3$, resulting in large aggregates. **b,** Cryo-EM micrograph of SAVED-CHAT incubated with $cA_3$ and vitrified immediately (within 10 seconds), preventing aggregation. **c,** Cryo-EM 2D classes of SAVED-CHAT-$cA_3$ filaments. **d,** Gold-standard Fourier Shell Correlation (FSC) of SAVED-CHAT-$cA_3$ 3D reconstruction. **e,** Directional FSC of reconstruction, corresponding to **d**. **f,** Euler orientation distribution plot. **g,** Map-to-model FSC, with a resolution of ~3.4 Å at the 0.5 threshold. **h,** SAVED-CHAT-$cA_3$ filament reconstruction coloured by local resolution.

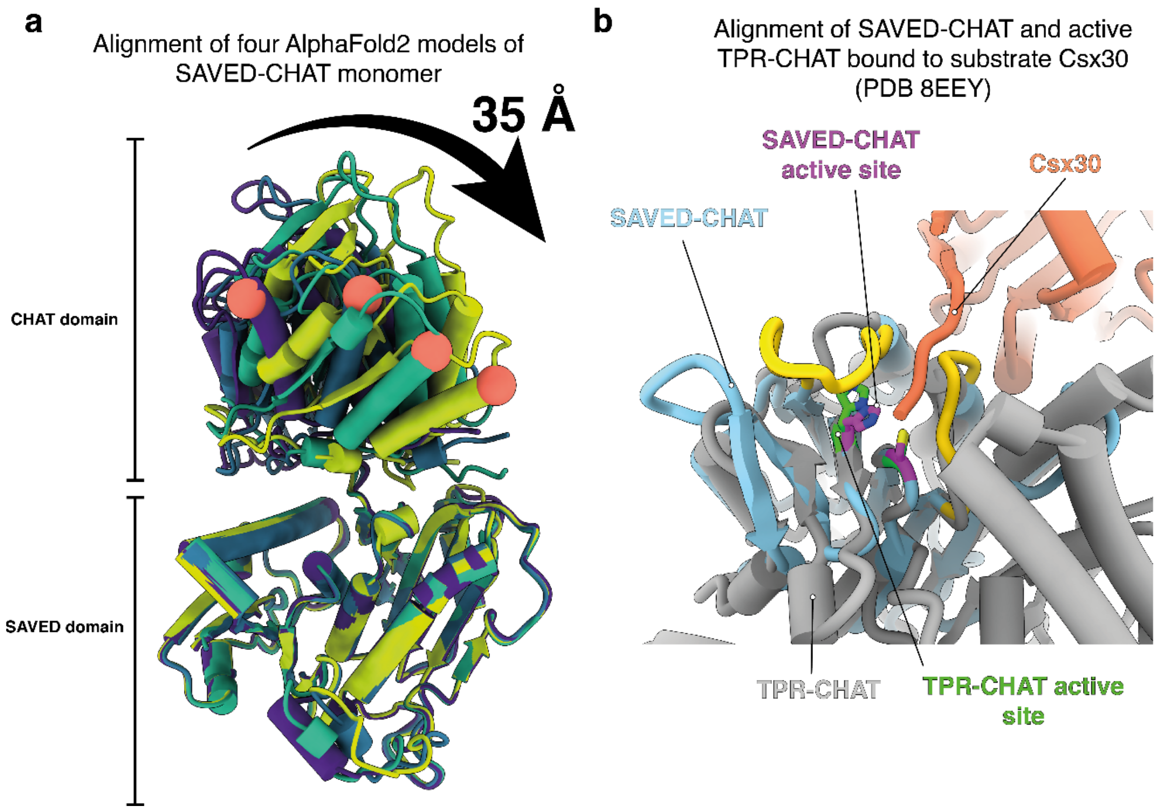


**Ext. Data Fig. 12. Structural analysis of SAVED-CHAT. a,** Flexibility of SAVED-CHAT monomer. Four AlphaFold2 models (colored purple, dark cyan, green and chartreuse) were aligned based on the SAVED domain. Red sphere corresponds to the same residue in all four models, highlighting the flexibility. **b,** Alignment of a single SAVED-CHAT proteolytic active site with TPR-CHAT (Csx29) bound to type III-E CRISPR effector in complex with activating non-self target RNA and substrate peptide Csx30. TPR-CHAT and SAVED-CHAT His-Cys active site catalytic dyad aligned in both structures, poised for cleavage of Csx30, confirming that the structure of SAVED-CHAT is in an active conformation.

**Ext. Data Table 1. Plasmids used in this study.**
(* will be added upon acceptance of the manuscript)

| Plasmid name | Main components | Addgene reference* |
|---|---|---|
| pJS-BCD-Strep-SAVED-CHAT | N-terminal Strep-tagged SAVED-CHAT | |
| pJS-BCD-Strep-dSAVED-CHAT | N-terminal Strep-tagged dSAVED-CHAT (H375A / C422A) | |
| pJS-BCD-Strep-PCaspase | N-terminal Strep-tagged PCaspase | |
| pJS-BCD-Strep-dPCaspase | N-terminal Strep-tagged dPCaspase (H79A / C146A) | |
| pJS-BCD-Strep-PCc-σ | N-terminal Strep-tagged PCc-σ | |
| pJS-BCD-Strep-PCi | N-terminal Strep-tagged PCi | |
| pHochTypeIII | *csb2*, *cmr1-cmr6*, minimal CRISPR array with one spacer | |
| pSAVED-CHAT_PCaspase_PCi_PCc-σ | SAVED-CHAT, PCaspase, PCi & PCc-σ | |
| pSAVED-CHAT | SAVED-CHAT | |
| pPCaspase | PCaspase | |
| pSAVED-CHAT_PCaspase | SAVED-CHAT & PCaspase | |
| pSAVED-CHAT_PCaspase_PCi | SAVED-CHAT, PCaspase & PCi | |
| pEcNucC | *E. coli* NucC | |
| pEmpty | N/A | |
| pTarget | Target sequence | |
| pNon-target | Non-target sequence | |
